# Isolation and Characterization of Porcine Endocardial Endothelial Cells

**DOI:** 10.1101/2022.12.04.518781

**Authors:** Kathleen N. Brown, Hong Kim T. Phan, Elysa L. Jui, Marci K. Kang, Jennifer P. Connell, Sundeep G. Keswani, K. Jane Grande-Allen

## Abstract

The heart contains six different types of endothelial cells, each with a unique function. We sought to characterize the endocardial endothelial cells (EECs), which line the chambers of the heart. EECs are relatively understudied, yet their dysregulation can lead to various cardiac pathologies. Due to the lack of commercial availability of this cell line, we developed a protocol for isolating EECs from porcine hearts and detailed our methodology for establishing populations of EECs through cell sorting. Additionally, we compared the EEC phenotype and fundamental behaviors to a well-studied endothelial cell line, human umbilical vein endothelial cells (HUVECs). The EECs were slightly smaller than HUVECs, and they stained positively for classic endothelial phenotypic markers such as CD31, von Willebrand Factor, and vascular endothelial (VE) cadherin. The EECs proliferated more quickly than HUVECs, yet migrated more slowly to cover a scratch wound assay. Finally, the EECs maintained their robust endothelial phenotype (expression of CD31) through more than a dozen passages. In contrast, the HUVECs showed significantly reduced CD31 expression in later passages. These important phenotypic differences between EECs and HUVECs highlight the need for researchers to utilize the most relevant cell lines when studying or modeling a disease of interest.

**Impact statement:** Many researchers model cardiovascular disease via tissue engineering to determine disease etiology on the cellular and molecular level. However, researchers usually rely on commercially-available cell lines, which can result in utilizing cells that are not specific to the region of the heart being modeled and could lead to incorrect conclusions. By providing a detailed protocol for isolating and purifying endocardial endothelial cells, we are enabling other researchers to access this cell line and model cardiovascular disease states more accurately.

## Introduction

Endothelial cells comprise 60% of the non-myocyte cellular composition of the heart.^1^ There are six endothelial phenotypes found within the heart: valvular, endocardial, coronary arterial, venous, capillary, and lymphatic.^2–4^ Each endothelial phenotype within the cardiovascular system serves a unique function and therefore differs in structure and behavior.^5^ For example, valvular endothelial cells cover the surface of the aortic, mitral, pulmonary, and tricuspid valves within the heart and are exposed to a high volume of circulating blood and mechanical forces during the cardiac cycle, whereas cardiac lymphatic endothelial cells are dispersed throughout the myocardium to drain fluids from the tissue and sprout vessels in response to injury.^1,3^ It has even been demonstrated that valvular endothelial cells on opposing sides of the aortic valve have different phenotypic signatures that can be detected.^6^ Furthermore, endothelial cells can profoundly shift their behavior, morphology, and cellular signaling in response to pathological environments.^4^ Since endothelial phenotype varies depending on location, and these variations may impact responses to pathological environments, the inclusion of region-specific endothelial cells in disease models can enhance understanding of the disease and increase understanding of disease mechanisms on the cellular level.

While the field of endothelial biology is continually expanding, many studies are performed used coronary, aortic, or human umbilical vein endothelial cells, with other endothelial phenotypes receiving relatively less investigation. One example is the endocardial endothelial cells (EECs), which line the chambers of the heart. EECs are often studied in development, as they form the inner layer of the heart tube during cardiac formation (as reviewed in ^7,8^). EECs demonstrate plasticity during cardiogenesis when they undergo endothelial-to-mesenchymal transition (EndMT) to contribute to heart valve formation.^9,10^ Postnatally, EECs are known to regulate oxygen and nutrient supply, mediate immune cell trafficking, and secrete vital signaling molecules to surrounding tissues.^8,11–14^ EECs play an essential role in cardiac formation and function, and dysfunction of EECs has been implicated in several cardiac diseases including congenital valve disease, hypoplastic left heart syndrome, and cardiac hypertrophy.^8,11,15–19^ Though there was initial speculation that EECs contribute to microvascular remodeling after myocardial infarctions, this was later disproven through genetic fate mapping, which demonstrated that mature EECs maintain their endocardial phenotype in this pathological environment ^20,21^.

We sought to characterize the EECs as a foundation for our investigations into the mechanisms of discrete subaortic stenosis (DSS).^22,23^ Since EECs serve as the barrier between circulating blood and the collagenous endocardium, just adjacent to the ventricular myocardium, this population of cells is exposed to a range of mechanical stimuli, including shear stresses due to cardiac blood flow. Unfortunately, it has been difficult to identify a commercial vendor for EECs. Therefore, to better understand the role of EECs in DSS and other cardiac diseases, we created a protocol to isolate EECs and thereby study their unique phenotype.

## Materials and Methods

### Preparation of Reagents and Workspace

The workspace and reagents were set up in advance of the heart dissection and enzymatic isolation of EECs. The following supplies were autoclaved: surgical tray, 600 mL beaker, two 1000 mL beakers, large surgical scissors, small curved surgical scissors, surgical hemostats, 12 cotton swabs, and two liters of phosphate buffered saline (PBS). Additionally, the following materials were gathered: spray bottle of 70% ethanol, six 8-ounce sterile specimen cups, 40 μm cell strainer, sterile towel drape. The dissociation solution contained collagenase and dispase. The stock collagenase solution was prepared with 25.2 mg of collagenase 2 (Worthington Biochemical) in 48.75 mL of Dulbecco’s Modification of Eagle’s Medium (Corning, #10-014-CV) and 1.25 mL of Penicillin-Streptomycin (Gibco, #15140122). The stock collagenase solution (0.05% collagenase) was then sterile filtered. The dissociation solution was prepared by combining 30 mL of the freshly prepared stock collagenase solution with 20 mL of dispase in Hank’s balanced salt solution (Stemcell Technologies, #07913), resulting in a 60% collagenase, 40% dispase solution. The tube was gently inverted to mix.

To prevent contamination, the heart dissection and isolation steps were performed in a biological safety cabinet (BSC), which required setting up the workspace in advance. The three sterile beakers were brought into the empty BSC. In the 600 mL beaker, 300 mL of 70% ethanol was added. The sterile surgical tools (scissors and hemostats) were then placed in the beaker with ethanol. Next, 800 mL of sterile PBS was added to both 1000 mL beakers. In one of the beakers, 40 mL of antibiotic-antimycotic (ABAM) (Gibco, #15240062) was added to create a 5% v/v solution. All the beakers containing the wash reagents remained at room temperature throughout the protocol. Finally, the sterile specimen cups, sterile drape, and sterile surgical tray were brought into the biological hood. The drape was opened and laid down first, then the surgical tray was placed on top. The lids of the specimen cups were removed so that the ventricles could easily be added to the cups.

### Heart Dissection

To obtain sufficient EECs, the isolation process required fresh tissue from multiple porcine hearts. For each EEC harvest, six porcine hearts were obtained from a local abattoir (Animal Technologies, Tyler, TX) and shipped overnight on wet ice. Fresh gloves were used to transport each heart from the packaging into the dissection workspace onto the surgical tray. The following procedure was completed for each heart individually while the other hearts remained on ice.

To obtain access to the left ventricle, the surrounding tissue was dissected off the heart, including the mitral valve. The large surgical scissors were used to remove the atria from the heart to expose the ventricles. The right ventricular wall tissue was then dissected away, leaving the left ventricle intact. The aorta was also removed by cutting immediately below the aortic valve. An incision was made on the exterior of the left ventricle immediately below the epicardial fat band, being careful to cut through the ventricle wall for the entire circumference of the ventricle while leaving the mitral valve intact. Once this was complete, the mitral valve was attached to the ventricle only by the chordae tendineae. Using the small surgical scissors, each chordal attachment of the mitral valve was carefully dissected from their attachment at the papillary muscle within the left ventricle.

After this dissection, the heart was placed into the beaker of PBS for a 5-minute rinse, then transferred using sterile hemostats to the PBS+ABAM for a second 5-minute rinse. After rinsing, the apex of the ventricle was placed into a sterile specimen cup, which supported the ventricle and held it upright. If the ventricle was too large to fit into the specimen cup, the exterior surface of the ventricle walls were trimmed down. The ventricle was then filled with PBS + ABAM, and set aside until all six hearts were similarly dissected and washed.

### EEC Dissociation and Purification

After preparing all the ventricles, they were filled with the dissociation solution. The PBS+ABAM solution was aspirated from each ventricle using a plastic aspirating tip, taking care to avoid puncturing the tissue. Next, each ventricle was filled with the dissociation solution, typically 7 ml of dissociation solution per ventricle but exact volume varied depending on ventricle size. The lids were then placed on the sterile cups, which were then transferred to a clean surgical tray and incubated at 37°C for 90 minutes.

After the incubation, the tray of ventricles was returned to the BSC. The dissociation solution was collected from each ventricle and transferred to a 50 mL conical tube fitted with a 40 μm cell strainer to filter out tissue debris. At this time, the dissociation solution was pooled from all six hearts. The inner surface of each left ventricle was swabbed three times using two sterile cotton swabs per ventricle to collect any remaining endothelial cells. Each cotton swab was then held over the 50 mL collection tube and washed with 1 mL of fresh dissociation solution to remove the cells from the swab. This step was repeated for each ventricle.

Once all of the dissociation solution was collected, the tube was centrifuged for 5 minutes at 150 x*g* to obtain a cell pellet. The supernatant was aspirated, and the cell pellet was resuspended in 10 mL of Endothelial Cell Growth Medium-2 (EGM-2, Lonza). The cell suspension was plated into a 75 cm^2^ tissue culture treated flask and cultured in a 5% CO_2_ incubator at 37°C. The growth medium was changed every other day.

To purify the EECs from any other cells that may have been harvested during the process, the cells were sorted at ~75%confluency. A magnetic bead sorting kit (CELLection Pan Mouse IgG Kit, Invitrogen) was used according to manufacturer’s instructions. The antibody used to coat the beads and sort the cells was PECAM-1 [CD31] Monoclonal Antibody (EMD Millipore, #MAB1393). The EECs were bead-sorted at passage 1 and passage 2 to ensure cell purity. Three populations of EECs were isolated and sorted before freezing and storing in liquid nitrogen until characterization and testing. The freezing media used to store the cells in was 80% EGM-2 media,10% dimethyl sulfoxide (DMSO, Sigma-Aldrich), and 10% fetal bovine serum (FBS, Fisher Scientific).

### HUVECs Cell Culture

We sought to compare the phenotype of the primary EEC cell line to a standard endothelial cell line: human umbilical vein endothelial cells (HUVECs). The HUVECs were obtained from Lonza (#C2519A). Once received, the HUVECs were seeded into a 75 cm^2^ tissue culture treated flask with 10 mL of EGM-2 and cultured in a 5% CO_2_ incubator at 37°C. The growth medium was changed every other day. After expanding the HUVECs, the cells were frozen down and stored in liquid nitrogen until testing. The freezing media used to store the cells in was 80% EGM-2 media, 10% DMSO, and 10% FBS.

### Immunofluorescence Staining of EECs

We stained the EECs for common endothelial markers to visualize the proteins within the cells. To prepare for immunofluorescence staining, the EECs were seeded onto glass slides. Sterile glass slides were placed into 100 mm petri dishes and coated with a 5% w/v gelatin in PBS solution for 10 minutes. The gelatin solution was aspirated off the glass slide and the surrounding petri dish. EECs were seeded onto the gelatin-coated glass slides at a density of 0.8 x 10^6^ cells per dish. The petri dish was then filled with 12 mL of EGM-2 media and incubated at 37°C. The media on the petri dish was changed every other day until the cells on the slide reached confluency.

Once confluent, the glass slide of EECs was fixed with 4% paraformaldehyde at room temperature for 10 minutes. The samples were washed with PBS three times for 5 minutes each. To permeabilize the cells, 0.2% Triton X-100 was added to the samples for 15 minutes, and again washed with PBS three times for 5 minutes each. Then the samples were blocked with a solution of 1% w/v of bovine serum albumin (BSA) in PBS at room temperature for one hour. After blocking, the samples were incubated with the primary antibody solutions overnight at 4°C (suspended in 1% BSA in PBS, see **Table 1** for antibody ratios). The next day, the samples were washed with PBS three times for 5 minutes each, then incubated with the secondary antibody solutions (suspended in PBS, see **Table 1** for antibody ratios) at room temperature for 1 hour, while protected from light. For samples stained with phalloidin (Thermo Fisher, #A34055, 1:100 dilution), the stain was added to the samples after blocking and incubated with the stain for 1 hour at room temperature. The samples were again washed with PBS three times for 5 minutes each. Finally, two drops of ProLong Glass Antifade Mountant with NucBlue (Thermo Fisher) were added to each slide of cells, and then a coverslip was added. Air bubbles were pushed out of the mounting media utilizing a 200 μL pipette tip. The samples were cured at room temperature and protected from light for 12 hours before imaging on Nikon Ti2 eclipse and Nikon A1 confocal microscopes.

**Table 1.** Primary and secondary antibody ratios used for immunofluorescence staining.

| **Primary Antibody** | **Primary Antibody Ratio** | **Corresponding Secondary Antibody** | **Secondary Antibody Ratio** |
| --- | --- | --- | --- |
| Mouse PECAM-1 [CD31] (EMD Millipore, #MAB1393) | 1:100 | Goat anti-Mouse IgG, AlexaFluor 555 (Invitrogen, #A-21422) | 1:100 |
| Rabbit pAb to von Willebrand Factor (Abcam, #ab6994) | 1:100 | Goat anti-Rabbit IgG, AlexaFluor 532 (Thermo Fisher, #A-111009) | 1:100 |
| Rabbit pAb to VE Cadherin (Abcam, #ab33168) | 1:500 | Goat anti-Rabbit IgG, AlexaFluor 532 (Thermo Fisher, #A-111009) | 1:100 |

### Scratch Wound Migration Assay

To assess cell migration rates, EECs and HUVECs were seeded into 6-well tissue-culture-treated plates at 0.3 x10^6^ per well. The cells were cultured with EGM-2 media until confluent. To begin the scratch wound assay, the media was aspirated from each sample, and the tip of a 1000 μL pipette tip was used to scratch a wound into the center of each well. Fresh media was added to each well, the plate was taken immediately for imaging on a Nikon T2 eclipse microscope to obtain brightfield images of the 0-hour time point for each sample. The samples were then imaged at 4 hours, 8 hours, and 24 hours after the scratch was made. The images were analyzed in ImageJ using the wound healing size tool plugin created by Suarez-Arnedo ^24^, which detects the border of the wound in each image and calculated the size of the wound or blank space on the sample. By evaluating the size of the wound over time, this analysis allowed us to compare the migration rates of EECs and HUVECs.

### Cell Proliferation Assay

To track cell proliferation over time, HUVECs and EECs were seeded into 96-well plates at 5 x10^3^ cells per well. The cells were cultured with EGM-2 media and incubated for 0, 24, 48, 72, and 96 hours. At each time point, the media was removed from the wells and the cells were incubated in 2 μM calcein AM in PBS (Live/Dead Assay, Thermo Fisher) for 30 minutes, then imaged immediately on a Nikon T2 eclipse microscope. The images were processed in ImageJ and the particles in the live channel were analyzed and quantified.

### Flow Cytometry

In preparation for flow cytometry, EECs and HUVECs were seeded into 75 cm^2^ tissue culture treated flasks at a density of 2.1 x10^6^ cells per flask. The cells were cultured with EGM-2 media. The cells were lifted and assessed by flow cytometry at ~75% confluence.

To lift the cells, the flasks were rinsed with PBS and then 10 mL of Accutase (Sigma-Aldrich) was added to each flask and incubated for 10 minutes at 37°C. The flasks were then gently tapped to detach the cells. The Accutase solution was transferred to a conical tube and centrifuged for 5 minutes at 150 x*g* to obtain a cell pellet. The cells were then resuspended in 1% FBS in PBS at a density of 1 x10^6^ cells per 100 μL. At this point, the cells were divided into multiple microcentrifuge tubes for testing. The CD31 antibody conjugated to fluorescent molecule APC (Thermo Scientific, #MA5-16769, 10 μL) or the APC isotype control (Thermo Scientific, #17-4714-42, 5 μL) was added to their respective samples tubes at concentrations recommended by the manufacturer. The samples were protected from light and incubated with the antibody at room temperature for 15 minutes. After incubation, 1 mL of 1% FBS in PBS was added to each tube for washing. The samples were centrifuged to obtain a cell pellet, then the supernatant was removed using a micropipette tip. The samples were washed a second time, and the cells were resuspended in ~500 μL 1% FBS in PBS and transferred to polystyrene 5 ml flow cytometry tubes (MTC Bio). The samples were placed on ice and protected from light until flow cytometry could be performed. Flow cytometry was performed on a Sony MA900 flow cytometer, and analysis was performed in FlowJo Version 7.

### Statistics

Statistical analysis was performed in GraphPad Prism Version 9.4.1. A one-way ANOVA test was performed on the data, with post-hoc analysis performed using Tukey’s test. Data in Figures 4 and 5 were graphed as means ± standard deviation (SD). Data in Figure 7 was graphed as individual data points. Results with p-value < 0.05 were considered statistically significant.

**Figure 1.**
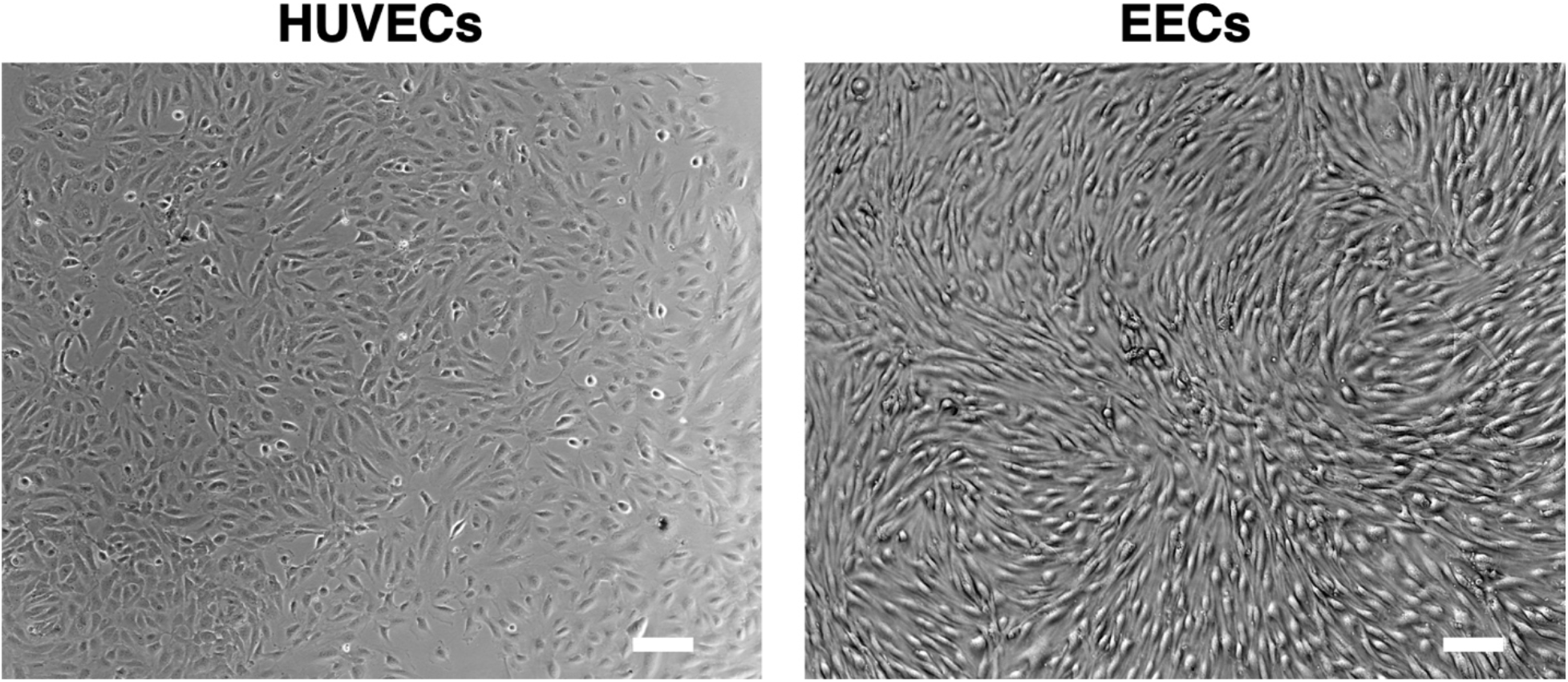
Brightfield images of human umbilical vein endothelial cells (HUVECs) and endocardial endothelial cells (EECs). Scale bar = 2000 μm.

**Figure 2.**
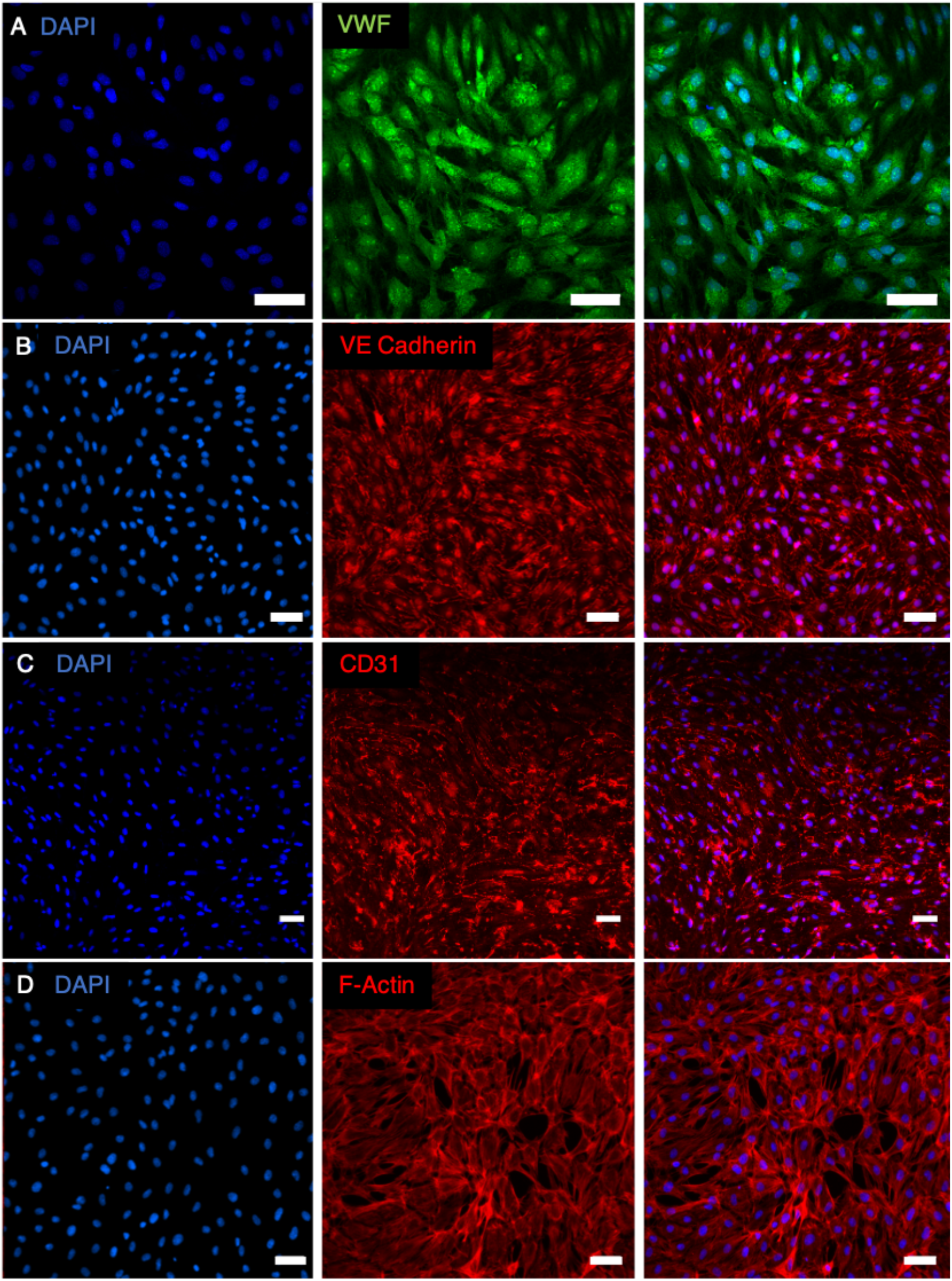
Immunofluorescence staining of EECs. In Row A, the EECs were stained for for von Willbrand Factor. In Row B, the EECs were stained for VE Cadherin. In Row C, the EECs were stained for CD31. In Row D, the EECs were stained for F-actin. Scale bar in Rows A, C, and D = 50 μm. Scale bar in Row B = 100 μm.

**Figure 3.**
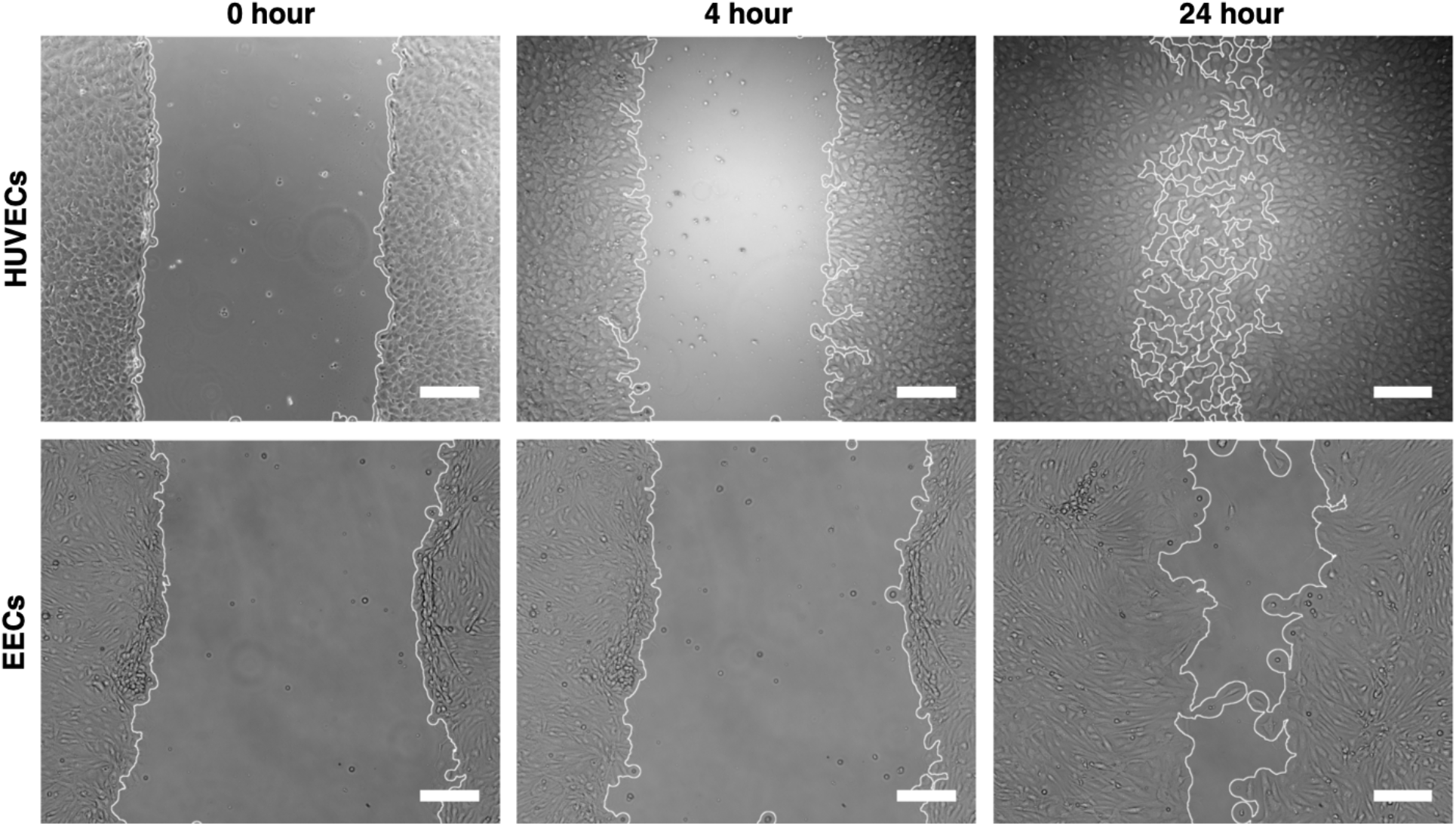
Brightfield images of HUVECs (top row) and EECs (bottom row) during the scratch wound migration assay at 0 hours, 4 hours, and 24 hours. The images shown also demonstrate the capability of the ImageJ wound healing size tool plugin, which oultined the edges of the wound created and calculated the size of the wound. Scale bar = 200 μm.

**Figure 4.**
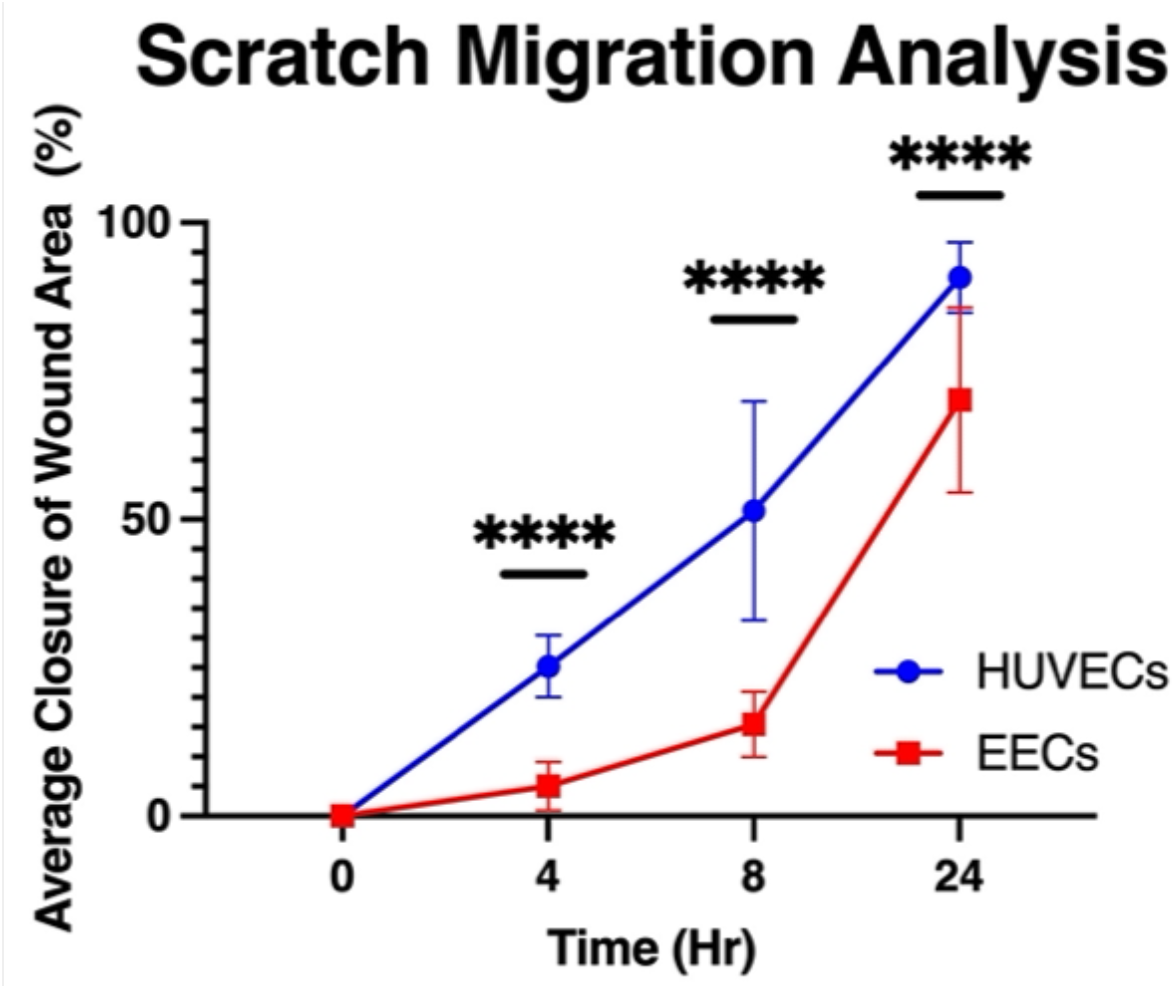
Graph comparing the migration rates of HUVECs and EECs at 0, 4, 8, and 24 hours. n=10. **** p < 0.0001.

**Figure 5.**
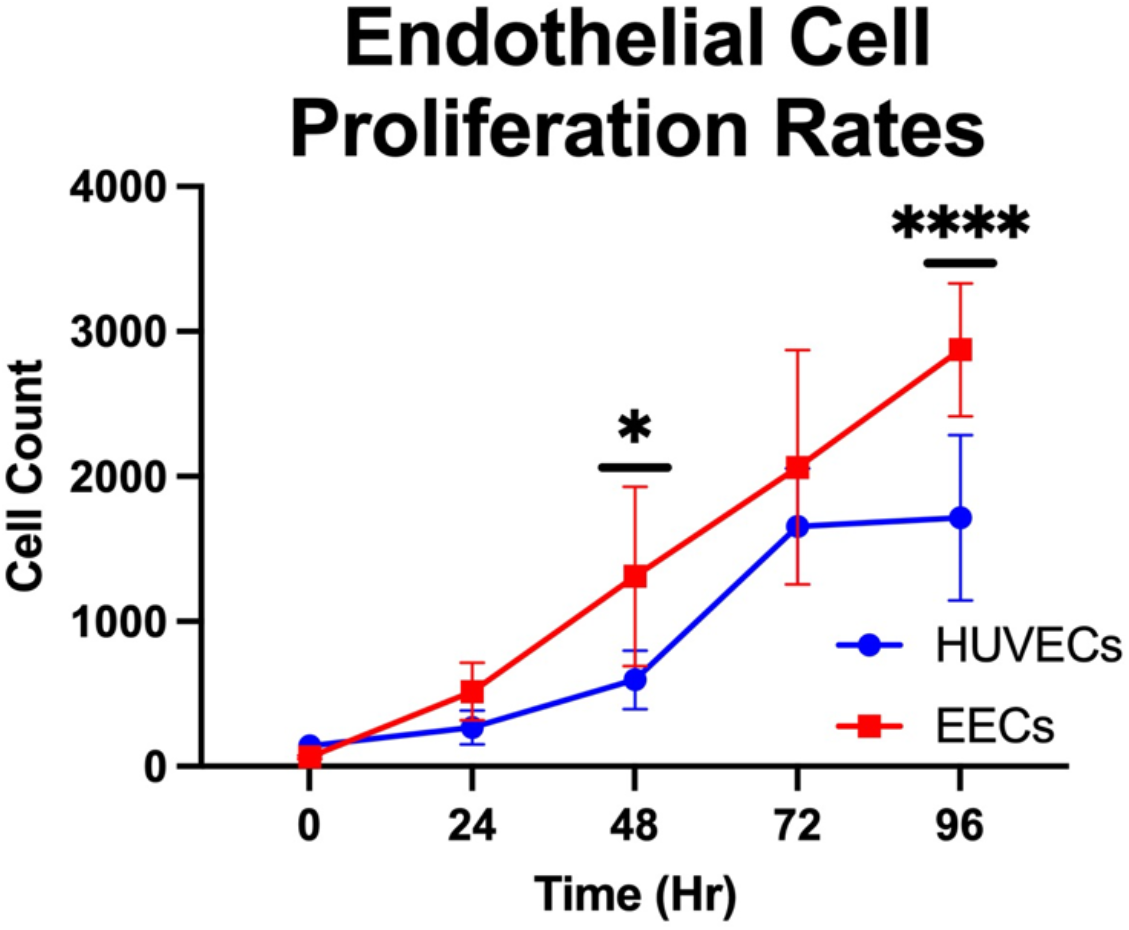
Graph comparing the proliferation rates of HUVECs and EECs calculated from live cell staining at 0, 24, 48, 72, and 96 hours. n=9. * p between 0.01-0.05; **** p < 0.0001.

**Figure 6.**
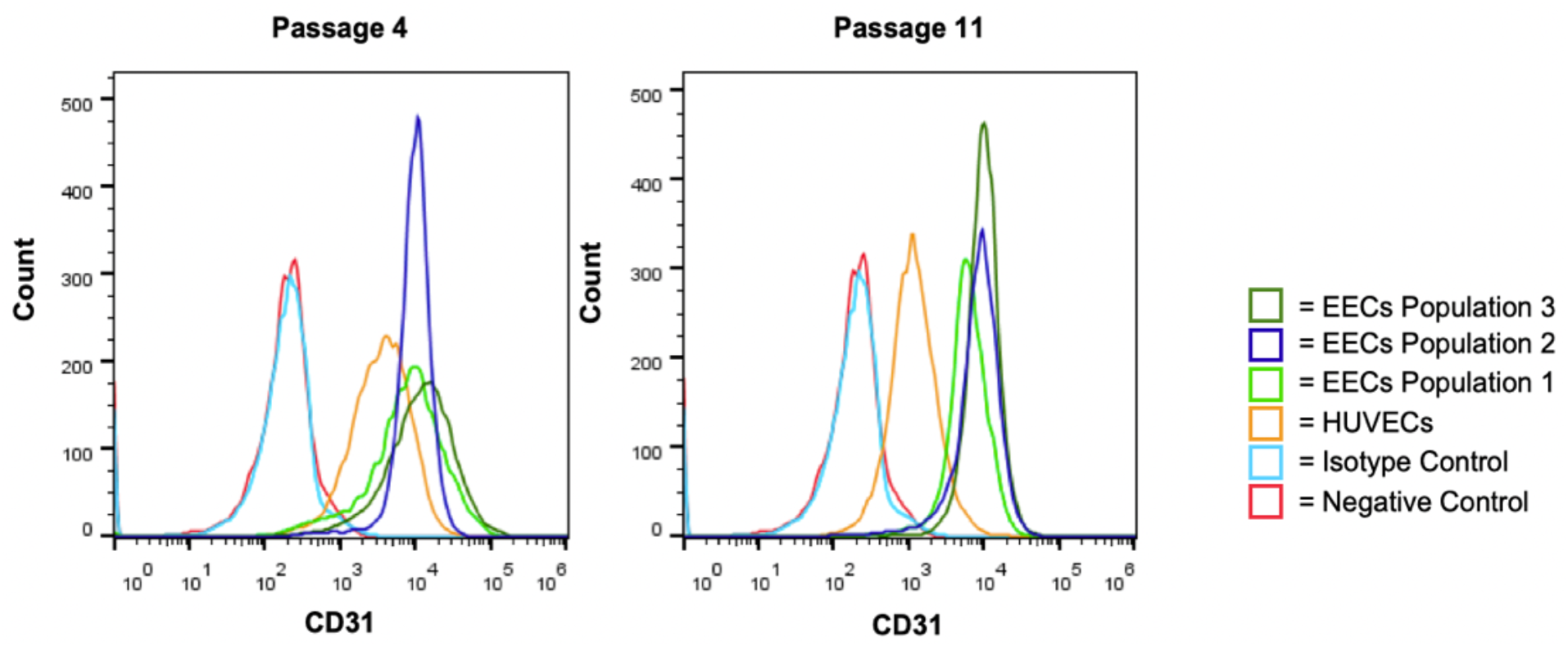
Flow cytometry results evaluating protein expression of CD31 betwen three populations of EECs, and a population of HUVECs across passages. Expression of CD31 across the samples can be seen at Passage 4 (graph on left) and Passage 11 (graph on right).

**Figure 7.**
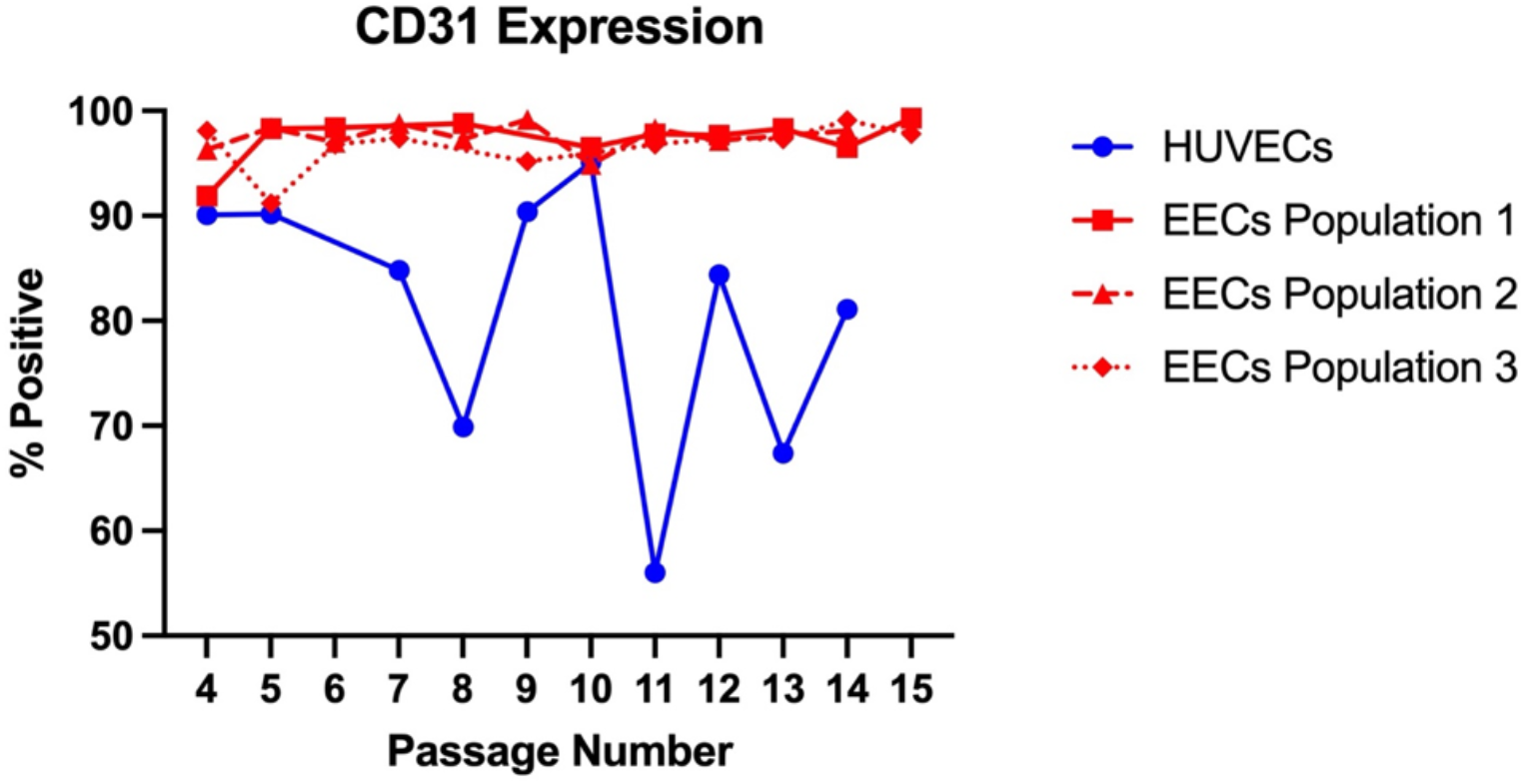
Graph of percentage of cells in each sample that positively expressed CD31 in the flow cytometry analysis from passage 4 through passage 15. Graph compares the HUVECs population to the three populations of EECs. # p between 0.0001-0.001 vs. all other groups.

## Experiment

### Phenotypic Characterization

Cellular phenotype is often evaluated by assessment of cellular morphology and expression of phenotypic markers through immunofluorescent staining.^4,25,26^ We aimed to compare the morphology to of EECs to HUVECs and assess differences observed from brightfield microscope images. Additionally, we evaluated the EECs to understand which endothelial phenotypic markers they expressed through immunofluorescent staining.

As demonstrated in brightfield images (**Figure 1**), the HUVECs cell line had a round and cobblestone morphology, which is characteristic of endothelial cells ^4,26^. The EECs had a round and cobblestone morphology, however there was elongation and swirling observed in the growth pattern as well. When comparing EECs to HUVECs, the cells were smaller and more elongated in their morphology.

Immunofluorescent staining was performed across the three populations of isolated EECs; representative images are shown in **Figure 2**. The EECs were stained for von Willebrand Factor (vWF), a protein that is only expressed in endothelial cells. vWF is contained within the Weibel-Palade bodies, which are storage granules that fuse in response to vessel injury to secrete vWF to initiate blood coagulation.^4,11,27–30^ Therefore, vWF typically presents as small granules within cells. This description is consistent with the staining shown in Figure 2A, indicating the cells positively expressed von Willebrand Factor. The EECs were also stained for vascular endothelial-cadherin (VE-Cadherin), a protein that is expressed on the membrane or surface of endothelial cells.^10,26,28,29,31–33^ In Figure 2B, the EECs positively expressed VE-Cadherin at the cellcell junctions. In Figure 2C, the EECs positively expressed CD31 on the membrane of the cells. In Figure 2D, the cells were stained with phalloidin, which selectively labels Factin filaments that compose the cytoskeleton.^12,26,27,31^ The F-actin stain demonstrates the distribution of actin filaments throughout the cells. Thus, the EECs positively expressed essential endothelial markers: CD31, von Willebrand Factor, and VE-Cadherin, which confirms their endothelial phenotype.

### Endothelial Behavior

Endothelial cell migration rates are an important feature *in vivo*, as endothelial cells participate in wound healing processes throughout the vasculature ^26^. Therefore, we compared the migration rates of EECs and HUVECs *in vitro* using the scratch wound assay, which was quantified by the ImageJ plugin. This assay demonstrated that the migration of HUVECs and EECs progressed at different rates (**Figure 3**). The HUVECs migrated faster than the EECs to close the wound created at all time points evaluated (**Figure 4**).

When evaluating migration rates of the cells, we considered how proliferation of the cells could affect the result of the migration assay. Therefore, we evaluated the proliferation rates of HUVECs and EECs. As shown in **Figure 5**, we found that the HUVECs proliferated more slowly than the EECs, with significantly fewer cells at the 48- and 96-hour timepoints. This result confirmed that the migration rate assay performed evaluated migration accurately, and proliferation did not influence the outcome of the migration assay results. Therefore, we concluded that HUVECs migrated faster than EECs.

### Phenotype Maintenance through Passaging

Cell surface markers are another important factor of cell phenotype. Many cell types, especially primary cells, change phenotype after being cultured and passaged for long periods. We sought to assess how well EECs maintain their endothelial phenotype through passages, and to compare their performance to HUVECs by evaluating CD31 protein expression with flow cytometry. The CD31 expression profile of the EECs and HUVECs shifted between passage 4 and passage 11, as shown in the flow cytometry data of **Figure 6**. The three populations of EECs maintained high expression levels of CD31 across all passages, however, the HUVECs expressed less CD31 at passage 11 than at Passage 4. In Figure 7, the percentage of cells positively expressing CD31 is graphed for three populations of EECs and one population of HUVECs. Each data point is the percent of CD31-positive cells for a passage; this was done for passages 4 to 15 of each population of cells. When comparing the CD31 expression across the three populations of EECs, there was no significant difference between the populations. When comparing the CD31 expression of the HUVECs to each EEC population, expression was significantly reduced, especially at later passages. Thus, the EECs maintained their endothelial phenotype across high passage numbers, particularly when compared to HUVECs.

## Discussion

Despite the importance of EECs in development and in disease pathologies such as myocardial infarctions, atrial fibrillation, and thrombosis ^11,18,19^, this cell line is largely understudied in the context of mature cardiac endothelium in physiological and pathological disease states. While this is an exciting potential avenue for research, there are significant challenges. There are currently no EEC lines commercially available, and thus EECs must be isolated by researchers from fresh cardiac tissue. Therefore, a thorough protocol on how to isolate and sort these cells has been greatly needed. Here, we have described a detailed procedure for the preparation and culture of EECs from porcine hearts. Importantly, we have confirmed the endothelial phenotype of EECs and demonstrated that this phenotype is stable over more than 10 passages, unlike HUVECs. Additionally, we have shown that the migration rate and proliferation rate are different from that of the more commonly studied HUVECs.

With the step-by-step guidelines provided here, investigators can isolate and purify EECs for a range of studies. It should be emphasized that to obtain sufficient numbers of EECs for experimentation at low passage numbers, this isolation procedure pooled together the cells from 6 porcine hearts at one time. Therefore, there remain challenges in isolating large numbers of EECs from human hearts, as they are a precious resource from suppliers such as the National Research Disease Interchange and cannot be obtained in such high quantities. Nonetheless, it will now be possible to incorporate EECs readily into disease models for cardiac pathologies for congenital and adult cardiac disease states. Evaluating the response of EECs to pathological environments such as those found in DSS has not yet been characterized; such studies could provide new insights into the progression and treatment of this and other conditions. Moreover, it is important to include EECs in models of cardiac diseases that are related to the cardiac lining, as how these cells respond would be more representative of the *in vivo* condition than one would find when modeling the disease using another endothelial cell line.^34^ While aortic endothelial cells are commercially available and easy to obtain, recent studies have found that EECs have a unique proteomic signature, even in comparison to aortic endothelial cells, further demonstrating the necessity of including EECs in cardiac disease models.^6,14^

This study also showed that EECs display the classical endothelial phenotypic characteristics, and that this phenotype is stable up to at least passage 15. EECs stained positively for typical markers of endothelial cells, but they were smaller and had a more elongated morphology than reported for other common endothelial cell lines.^4,26^ Importantly, we found that EECs maintained higher endothelial integrity over many passages than did HUVECs, a novel insight that could influence the choice of cell line for future research. HUVECs are often used as a model endothelial cell line as they are easy to obtain from a variety of suppliers and generally demonstrate the morphological and phenotypical signatures that are common to endothelial cells. However, HUVECs are not as viable in high passages, and show a deteriorating endothelial integrity after passages 5-6.^26,28^ Additionally, HUVECs are immune naïve in comparison to adult endothelial cells, which have developed immunological capabilities as they have been exposed to the native postnatal immune system prior to isolation.^35^ These results highlight the pitfalls of using a common model endothelial cell line.

Furthermore, the EECs showed a faster proliferation rate and a slower migration rate than HUVECs, which underscores the need for researchers to utilize endothelial cell lines that are specific to the tissue region under investigation. Often, cell lines that are convenient to obtain are chosen for *in vitro* disease models, but as shown here, there are important phenotypic differences for each cell subtype that could influence study findings. In characterizing these porcine EECs, we have demonstrated that this cell line should be an excellent resource for researchers who desire to study cardiac endothelial cells across many passages, and who aspire to model aspects of disease observed in left ventricular outflow tract and other endocardial pathologies.

Although these results showing the phenotypic stability of EECs are compelling, there were some experimental contexts that should be considered. To begin, porcine hearts were used in lieu of human hearts. The hearts of young adult pigs (6-12 months old) are commonly used as an experimental model for human hearts due to their anatomical and structural similarities and convenient availability.^36–38^ Pigs and sheep are also often used in surgical large animal models of diseases of the heart, valves, and cardiovascular system.^39,40^ In the future, it will be important to determine how well porcine EECs represent the behavior of human EECs. In addition, the scratch wound migration assay was not conducted in the presence of a proliferation inhibitor. Because the EECs were found to proliferate more rapidly than the HUVECs, their migration rate would have been even slower were such an inhibitor used in this study. Nonetheless, these studies were able to demonstrate significant differences between EECs and HUVECs.

In conclusion, we successfully isolated, purified, and cultured endocardial endothelial cells, the cells that line the left ventricle and left ventricular outflow tract of the heart. We demonstrated the robust endothelial phenotype of EECs, as the cells maintained their endothelial protein marker expression even to passage 15. We showed that there are important phenotypical differences between EECs and HUVECs, which emphasizes the need for researchers to utilize specific cell lines relevant to the diseases under investigation.

## Author Contributions

Brown: conceptualization (lead), investigating, formal analysis, writing – original draft (lead). Phan: investigating. Jui: investigating, formal analysis (support). Kang: conceptualization (support). Connell: conceptualization (support), formal analysis (support), writing – review & editing. Keswani: funding acquisition, supervision. Grande-Allen: conceptualization (support), funding acquisition, supervision, writing – review & editing.

